# Unraveling human, rodent and snake *Kolmioviridae* replication to anticipate cross-species transmission

**DOI:** 10.1101/2023.05.17.541162

**Authors:** Pierre Khalfi, Zoé Denis, Giovanni Merolla, Carine Chavey, José Ursic-Bedoya, Lena Soppa, Leonora Szirovicza, Udo Hetzel, Jeremy Dufourt, Cedric Leyrat, Nora Goldmann, Kaku Goto, Eloi Verrier, Thomas F. Baumert, Dieter Glebe, Valérie Courgnaud, Damien Grégoire, Jussi Hepojoki, Karim Majzoub

## Abstract

The recent discovery of Hepatitis D (HDV)*-like* viruses across a wide range of taxa led to the establishment of the *Kolmioviridae* family. Recent studies suggest that kolmiovirids can be satellites of viruses other than Hepatitis B virus (HBV), challenging the strict HBV/HDV-association dogma. Studying whether kolmiovirids are able to replicate in any animal cell they enter is essential to assess their zoonotic potential. Here, we compared replication of three kolmiovirids: HDV, rodent (RDeV) and snake deltavirus (SDeV) *in vitro* and *in vivo*. We show that SDeV has the narrowest and RDeV the broadest host cell range. High resolution imaging of infected cells revealed nuclear viral hubs with a peculiar RNA-protein organization. Finally, *in vivo* hydrodynamic delivery of infectious clones showed that both HDV and RDeV, but not SDeV, efficiently replicate in mouse liver, forming massive nuclear viral hubs. Our comparative analysis lays the foundation for the discovery of specific host factors controlling *Kolmioviridae* host-shifting.

## Introduction

For over 40 years, Hepatitis D virus (HDV) was the only known member of the unassigned genus *Deltavirus*^1–3^. Originally discovered in a Hepatitis B virus (HBV) infected patient, HDV was later shown to be a satellite of HBV^4,5^, a major cause of liver disease and cancer. Recently, the coincidental identification of HDV*-like* elements, in bird cloaca^6^ and snake brains^7^, provided the first evidence that HDV is not the sole representative of deltaviruses. Since then, several independent meta-transcriptomic studies have identified HDV-*like* sequences in a variety of samples originating from bats, rats, deers, marmots, birds, frogs, fishes and insects^8–12^. These discoveries indicated that this new viral family is far more diverse and widespread amongst the animal kingdom than originally thought. Evidence that HDV*-like* viruses found in snakes^13,14^, rodents^9,11^ and birds^11^ are able to replicate and the recent identification of thousands of sequences similar to deltaviruses in metatranscriptomes^12,15^ led to the establishment of a novel realm, *Ribozyviria* with a single family, *Kolmioviridae*, that includes the genus *Deltavirus* as well as seven other novel genera of kolmiovirids^16^.

Because other kolmiovirids were only recently discovered, most of our knowledge of the biology of these agents stems from research on HDV^17^. HDV possesses a negative-stranded, circular and highly self-complementary RNA genome of ∼1700 nucleotides, making it the smallest known virus able to infect animal cells^17,18^. An estimated 15 to 20 million individuals worldwide are HBV-HDV co-infected, and chronic co-infection is considered the most severe form of viral hepatitis, often leading to advanced liver disease and cancer^19–22^. HDV highjacks HBV surface antigens (HBsAg) for infectious particle formation and enters human hepatocytes via the sodium-taurocholate co-transporting polypeptide (NTCP) receptor, which dictates its liver tropism^23,24^. Once HDV gains access to host cells, viral transcription and replication are mediated by cellular RNA polymerases, independently of HBV^25,26^. The HDV genome encodes a single protein, the hepatitis delta antigen (HDAg) which exists in two forms, the small (S-HDAg) and the large (L-HDAg) that differs from S-HDAg by 19 additional amino acids (AAs) at its C-terminal end^27^. The S-HDAg is essential for viral RNA replication^28^, while a farnesylated form of L-HDAg is involved in HDV assembly^29,30^.

Although newly discovered kolmiovirids share similar genome size and organization with HDV, they appear to differ from HDV in many aspects. For instance, they are not restricted to the liver of infected animals. Swiss snake colony virus 1 (SwSCV-1) (hereafter referred to as Snake deltavirus or SDeV) was detected in the spleen, kidney, lung and brain of infected boa constrictors^7^. Likewise, Tome’s spiny-rat virus 1 (TSRV-1) (hereafter referred to as Rodent deltavirus or RDeV) was detected in the kidney, lung, heart and small intestine of infected spiny rats^9^. Importantly, none of the novel kolmiovirids has been linked to a *Hepadnaviridae* (HBV family) co-infection so far. In fact, Reptarena- and hartmaniviruses, commonly found in captive constrictor snakes^31^, were shown to act as helper viruses of SDeV^13^. Furthermore, HDV was recently shown to form infectious particles with envelope glycoproteins different from HBsAg (e.g. *Flaviviridae, Rhabdoviridae*)^32^. These observations lend support to the idea that kolmiovirids can invade virtually many cell types, when packaged with the appropriate viral envelope. Interestingly, a recent study proposed that these viruses are capable of host-shifting between highly divergent species, suggesting that the contemporary association between HDV and HBV likely arose following zoonotic transmission from a yet undiscovered animal reservoir^10^.

The capacity of these satellite viruses to enter virtually many cell types coupled to their exclusive reliance on host factors for replication begs the question: Can kolmiovirids replicate in any cell type they access? Here, we try to address this question using HDV, RDeV and SDeV infectious clones to characterize their replication in a variety of animal cell lines and in an *in vivo* mouse model.

## Results

### Comparison of human, rodent and snake delta antigens (DAgs)

HDAg harbors seven previously mapped regions: three RNA-binding motifs (RBMs), a nuclear localization signal (NLS), a coiled-coil domain, a helix-loop-helix motif (HLH) and a proline- and glycine-rich (PGR) region at the C-terminus^33–36^ (Fig.1A and Fig.S1B). Furthermore, post-translational modification of three AA residues in the S-HDAg have been shown to be important for HDV replication: Arg-13 methylation^37^, Lys-72 acetylation^38^ and Ser-177 phosphorylation^39^.

**Figure 1:**
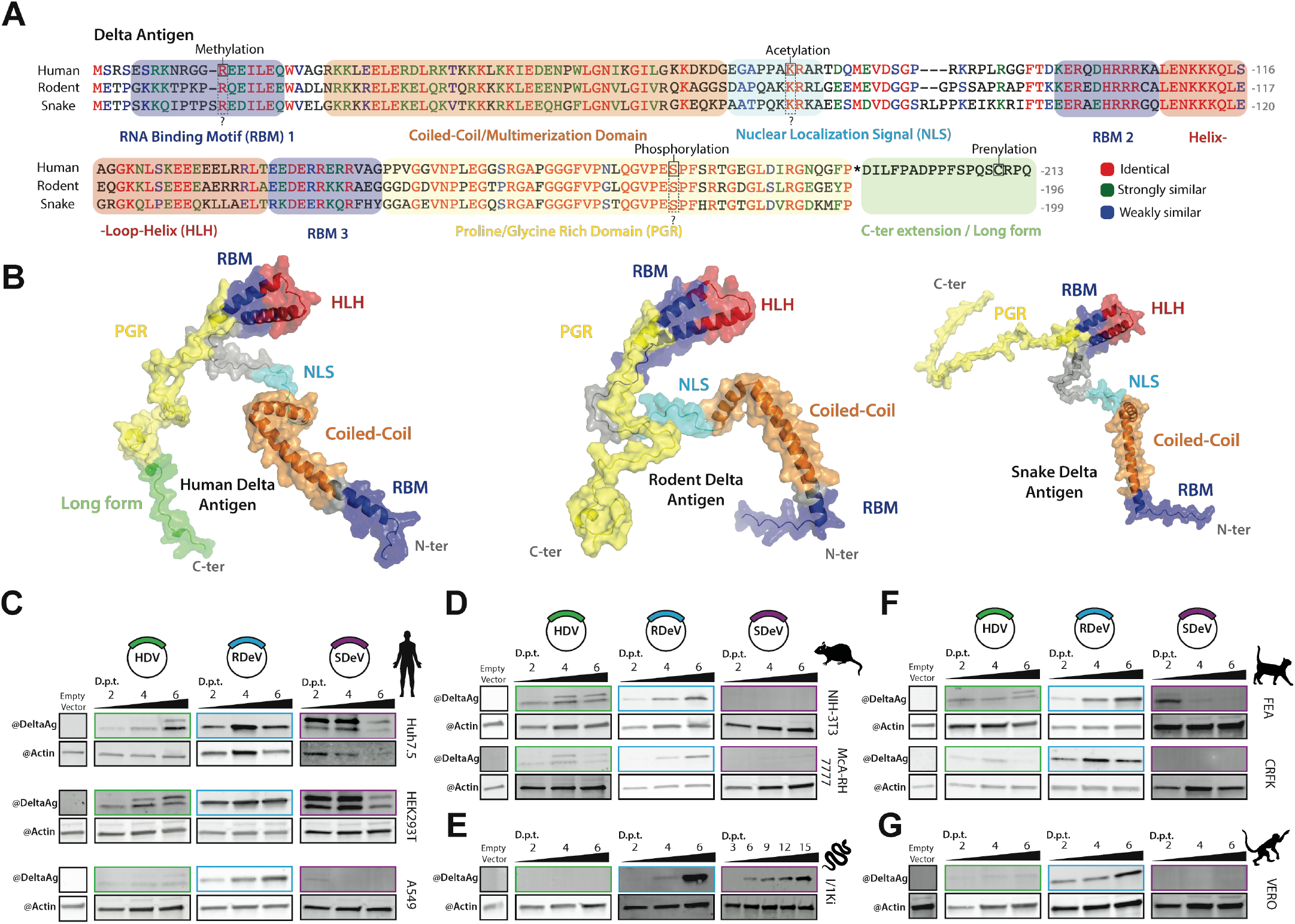
Comparison of HDV, RDeV and SDeV antigen’s amino acid sequences, predicted 3D structures and replication in immortalized cell lines. (A) Clustal Omega alignment of HDV, RDeV and SDeV amino acid sequences. Amino-acid are color coded by similarity, the characterized HDAg domains are indicated by the colored boxes and known post-translational modifications of HDAg are indicated by the dashed boxes. B) AlphaFold2 prediction of the 3D structure of human, rodent and snake delta antigens. The characterized HDV domains are indicated on all three antigens by color. (C-G) deltaviruses’ antigen accumulation in different immortalized cell lines. pcDNA3.1 plasmids encoding dimers of the human, rodent and snake deltavirus’ genome, or an empty backbone, were transfected in human (C), rodent (D), snake (E), cat (F) and monkey (G) cell lines. Cells were collected at 2,3,4,6,9,12 and 15 days post-transfection (d.p.t.) and delta antigen expression was analyzed by western blot. β-actin serves as a loading control.

We first sought to map known domains and modified residues present in HDAg, to conserved regions in rodent and snake DAgs, both on the primary AA sequence and on putative 3D structural models. To do so we performed: 1-a multiple AA alignment of HDAg (genotype 1 – HDV isolate Taylor), RDAg (RDeV isolate 183) and SDAg (SDeV isolate F18-5) (Fig.1A) from which we calculated a sequence conservation score (Fig.S1A-B), 2-an analysis of intrinsic disorder along the sequence of each DAg (Fig.S1B) and 3-3D structure predictions of each of the three antigens using AlphaFold2 (AF2), mapping different functional and structural motifs onto the structural models of HDAg, RDAg and SDAg monomers (Fig.1B, Fig.S1B). Overall, the analysis of sequence conservation showed that while HDAg and RDAg are most closely related, sharing 57% AA identity (with 69% similarity), SDAg is slightly more divergent with 49 and 56% AA identity to HDAg and RDAgs respectively, but equivalent in similarity (66 and 70% respectively) (Fig.S1A). The comparison of sequence conservation, structural disorder and AF2 confidence score profiles shows a conserved modular architecture of the DAgs (Fig.1B-S1B). The coiled-coil domain and HLH motif display strong sequence conservation and form stable structures that are predicted with high confidence by AF2 (70 to 90 pLDDT score) (Fig.1B-S1B). Based on the disorder score, the HLH motif appears less stable than the coiled-coil domain, and is flanked by two intrinsically disordered regions encompassing the NLS at its N-terminus and the PGR at its C-terminus. Interestingly, the NLS is the least conserved region between each DAgs, while the most conserved stretch of residues is in the PGR (>70% AA identity for AA 150-190 of HDAg) (Fig.S1B). Furthermore, the combination of high conservation, low disorder score and low AF2 confidence score in the PGR (Fig.S1B) suggests that this region may contain a short linear interaction motif that folds upon interaction with protein and/or RNA partners. Importantly, residues Arg-13^37^, Lys-72^38^ and Ser-177^39^ known to be post-translationally modified in HDAg, are conserved in both RDAg and SDAg (Fig.1A). Taken together, our analysis reveals a common modular architecture of DAgs, and the conservation of important structural and functional motifs, as well as posttranslationally modified AA residues previously identified in HDAg. These results suggest that similar functional motifs and post-translational modifications might govern kolmiovirid replication.

### Antibody cross-reactivity for detection of kolmiovirid DAgs

Kolmiovirid DAgs share sequence homology and therefore potential epitopes for antibody-based detection^9^. To allow reliable detection of DAgs, we tested different antisera for their cross-reactivity: six obtained from HDV-positive patients^40,41^ and one from a rabbit immunized with recombinant SDAg^7^. We cloned in mammalian expression vectors HDV, RDeV, SDeV, Chusan Island toad virus 1 (CITV-1), and dabbling duck virus 1 (DabDV-1) DAgs with a C-terminal FLAG-tag. Ectopic expression of these constructs in Huh7 cells, followed by immunoblotting, revealed that all tested antisera were able to cross-react and detect HDV, RDeV, SDeV and DabDV-1 DAgs (Fig.S1C). However, CITV-1 (toad) DAg was only detected by 2 out of the 7 tested antisera (Fig.S1C), in agreement with the phylogenetic divergence of this protein (Fig.S1D). Therefore, we can use available antisera to reliably detect kolmiovirid DAgs and thus compare and characterize the replication of RDeV and SDeV.

### HDV, RDeV and SDeV replication in human and animal cell lines

Although most studies have reported HDV replication in human hepatocytes, other kolmiovirids are not restricted to the liver of their animal hosts^7,9^. Moreover, HDV’s and SDeV’s capacity to form infectious particles using envelope proteins of various viruses^13,32^ allows kolmiovirid entry into virtually many cell types. However, it remains unclear if all cell types are permissive to kolmiovirid replication. To focus on replication, we bypassed the viral entry step and transfected established HDV, RDeV and SDeV infectious clones^9,13^ into a battery of immortalized animal cell lines, including cells of human, mouse, rat, snake, cat and monkey origins, and followed viral replication over time by immunoblotting (Fig.1C-G).

All tested human cell lines (Huh7.5 – liver derived, HEK293T – kidney derived and A549 – lung derived) supported HDV and RDeV replication as shown by the accumulation of DAg over time (Fig.1C), with HDV replicating very poorly in the A549 cell line. Interestingly, SDAg accumulated to detectable levels in HEK293T and Huh7.5 cells, peaking at 4 days post-transfection (d.p.t) and decreasing at 6 d.p.t. (Fig.1C). Intriguingly, we were able to detect two forms of the SDAg, with distinct sizes, in the HEK293T and Huh7.5 cells transfected with the SDeV clone (Fig.1C). Rodent cell lines (NIH-3T3 – mouse fibroblasts and MCA-RH 7777 – rat liver derived) supported HDV and RDeV but not SDeV replication (Fig.1D). The snake cell line I/1Ki (derived from *boa constrictor* kidney) supported RDeV and SDeV replication but was refractory to HDV (Fig.1E). Interestingly, unlike in human HEK293T and Huh7.5 cells (Fig.1C), but in agreement with Hetzel *et al*.^7^, only one form of the SDAg was detected in the I/1Ki cell line (Fig.1E). Two feline cell lines (FEA – cat embryonic fibroblasts and CRFK – kidney cortex derived) were able to support RDeV, but supported very poorly HDV and SDeV replication (Fig.1F). Vero cells (African green monkey kidney derived) only supported RDeV replication (Fig.1G). In conclusion, our data show varying abilities of different cell lines to support HDV, RDeV and SDeV replication. While all tested cell lines were permissive to RDeV replication, HDV replicated efficiently only in certain cell types (Fig.1C-G). SDeV replication appeared to be the most restricted, as only the snake I/1Ki cell line supported efficient replication over time (Fig.1E). Intriguingly, HEK293T and Huh7.5 cells supported SDeV replication to some extent, and two forms of the SDAg accumulated, a large and a small one (Fig.1C), unlike in the snake I/1Ki cell line, where only one form of the SDAg is produced^7^ (Fig.1E).

### Effect of ADAR-1 editing on HDV, RDeV and SDeV DAg production

During HDV replication, the synthesis of the S- and L-HDAgs, is due to the translation of two distinct viral mRNAs^42^. An ADAR-1-dependent editing event of the HDV antigenomic RNA, transforms the amber stop codon (UAG) of the S-HDAg into a tryptophan (W) codon (UGG), extending the reading frame by 19 codons/AAs, thus allowing the transcription of a distinct mRNA coding for the L-HDAg (Fig.2A)^43–46^. SDeV genome also possesses an amber stop codon that could be potentially edited into a W (UGG) in the end of the small SDAg (S-SDAg), giving rise to the putative larger SDAg (L-SDAg) form, extended by 22 AAs relative to the smaller one (Fig.2A). We thus sought to investigate if the two SDAgs forms observed in human cells (Fig.1C), were due to an editing event of SDeV RNA, leading to the production of two forms of DAg^42–46^.

**Figure 2:**
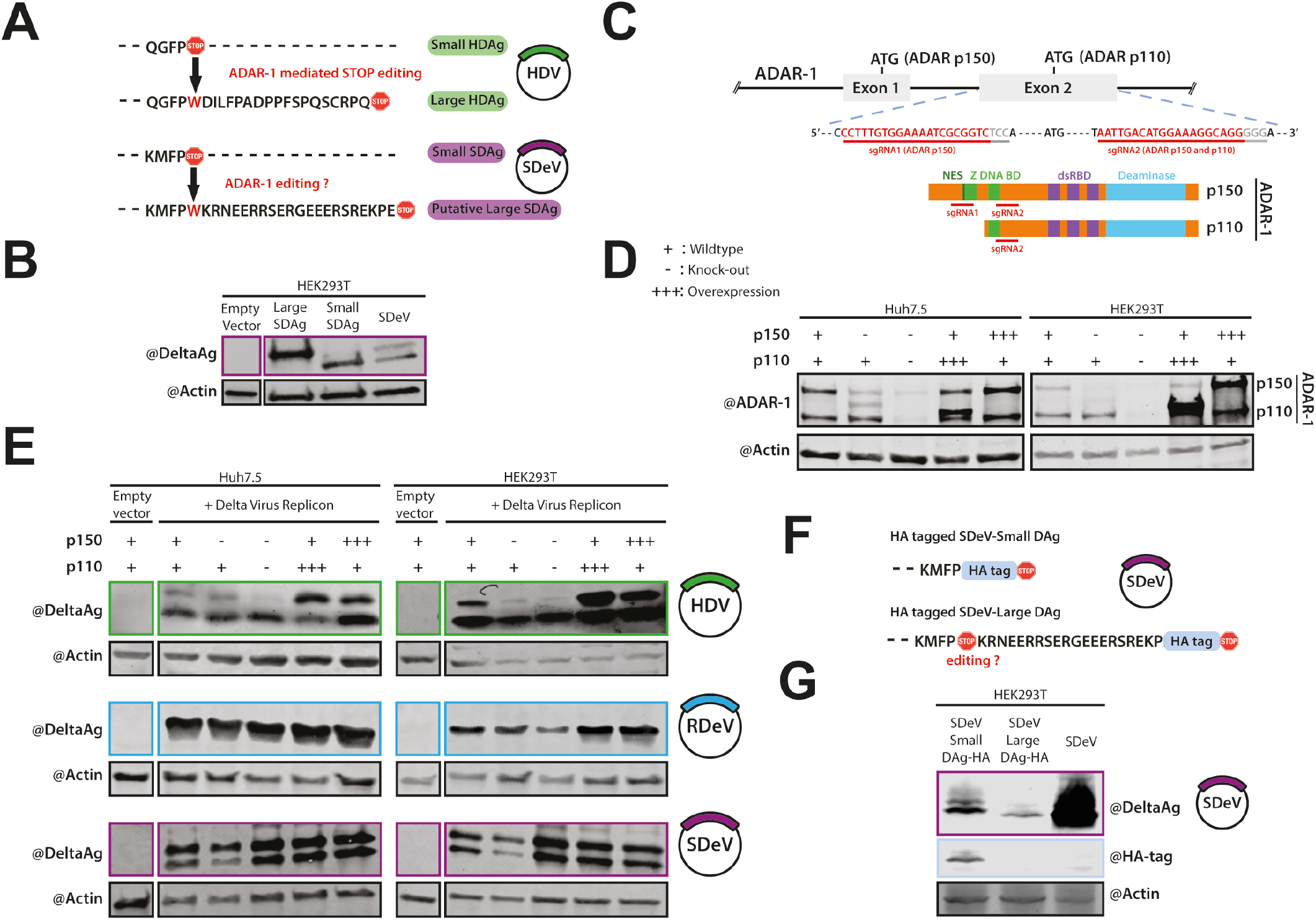
Editing and DAgs accumulation. A) Western blot analysis of SDAgs forms. Cells were collected 4 days post-transfection with plasmids encoding small or putative large snake delta antigens and protein extracts were analyzed by western blot probing DAgs and β-actin expression. B) Schematic representation of the C-terminal amino acid sequences of the small and large HDAg resulting from ADAR-1 mediated editing of HDV (*Upper*), and the small and putative large SDAg resulting from putative editing of SDeV (*Lower*). C) Schematic representation of the ADAR-1 locus targeted by the different sgRNAs for KO generation (*Upper*) and the ADAR-1 isoforms, p110 and p150, including the functional domains for each isoform and the localization of the sgRNAs targeted regions (*Lower*). D) Western blot analysis of WT, ADAR-1 KO or overexpressing cells lines. Protein extracts from ADAR-1 overexpressing (p110 or p150), ADAR-1 KO (p110 and p150 or p150 only) and control Huh7.5 and HEK293T cell lines were analyzed by western blot to detect ADAR-1 expression and β-actin E) Western blot analysis of the human (*Upper*), rodent (*Middle*) and snake (*Lower*) DAg forms in ADAR-1 overexpressing (p110 or p150), ADAR-1 KO (p110 and p150 or p150 only) and control Huh7.5 (*Left panels*) and HEK293T (*Right panels*) cell lines. Cells were transfected with an empty pcDNA3.1 plasmid or with pcDNA3.1 plasmids encoding dimers of the HDV, SDeV or RDeV genomes and collected 9 days post-transfection (d.p.t.) for HDV transfected Huh7.5 cells, 6 d.p.t for HDV transfected HEK293T cells, 6 d.p.t. for RDeV transfected Huh7.5 and HEK293T cells and 3 d.p.t. for SDeV transfected HEK293T and Huh7.5 cells. Protein extracts were analyzed by western blot for DAg and β-actin expression. F) Schematic representation of the C-terminal amino acid sequences of the HA tagged small and putative large SDAgs resulting from a putative ADAR-1 mediated editing of SDeV RNA. G) Western blot analysis of HA tagged SDeV antigen expression. HEK293T cells were transfected with 1.2x SDeV genome pCAGGS vector harboring an HA tag after the SDeV small antigen, the SDeV putative large antigen, or not harboring any HA tag. Cells were collected 2 days post-transfection and protein extracts were analyzed by western blot for DAg, HA tags or β-actin expression.

To verify if the two SDAg forms observed in human cells (Fig.1C) correspond to S-SDAg and L-SDAg (Fig.2A), we cloned their respective open reading frames into a mammalian expression vector. Transfection of these constructs in HEK293T cells followed by immunoblotting side-by-side with SDeV infected cell lysates (Fig.2B) showed that the L-SDAg migrates to a similar position to that of the higher molecular weight form of SDAg observed during SDeV replication (Fig.2B), suggesting that the two observed forms could well be the result of ADAR-1 editing.

To test this hypothesis, we knocked-out (KO) both ADAR-1 forms (p110 and p150) by CRISPR-Cas9^47^ (Fig.2C-D) or overexpressed both forms in two different human cell lines and verified SDAg production (Fig.2D). As expected, ADAR-1 KO cell lines showed compromised L-HDAg production, while ADAR-1 overexpression enhanced L-HDAg production (Fig.2E). To our surprise, neither KO nor overexpression of ADAR-1 affected RDAg or SDAg expression (Fig.2E), suggesting that the two SDAg forms are produced thanks to an ADAR-1-independent mechanism.

To study if the two SDAg forms are due to editing by another cellular factor (e.g. ADAR-2), we engineered SDeV infectious clones with a hemagglutinin (HA)-tag (Fig.2F). In the first construct, the HA-tag is adjacent to the C-terminus of the small SDAg (S-SDAg clone) and in the second, the HA-tag is adjacent to the 21 putative extra AAs, translated only if an editing event occurs (L-SDAg clone) (Fig.2F). Immunoblot of HEK293T cells transfected with these constructs shows that while both can replicate (Fig.2G), only the S-SDAg clone could be detected by an anti-HA antibody (Fig.2G), indicating that the stop codon editing does not occur in SDeV. Interestingly, the S-SDAg clone produced a doublet band detectable also with anti-HA antibody, suggesting that the larger SDAg species observed during SDeV replication in human cells might be due to post-translational modification(s).

### HDV, RDeV and SDeV RNA and DAg accumulation patterns in human, rodent and snake cells

More than a decade ago, several studies have shown that both HDAg and HDV RNAs localize to nuclei of infected cells where viral replication takes place^28,48–52^. To characterize and compare HDV, RDeV and SDeV RNA and DAg localization patterns in infected cells, we first generated persistently infected clonal cell lines for each virus. Fluorescence-activated cell sorting (FACS) served to select and amplify single Huh7.5.1, NIH-3T3 and I/1Ki cell clones, transfected with HDV, RDeV and SDeV infectious clones respectively. RT-qPCR and western blot was used to identify several positive clones for each virus/cell line combination.

Immunofluorescence (IF) imaging using IgGs purified from HDV-positive patients’ sera revealed a nuclear localization of HDAg, RDAg and SDAg in human, rodent and snake cells respectively (Fig.3A). We then investigated if HDV, RDeV and SDeV replication causes the formation of double-stranded RNA (dsRNA), a replication intermediate hallmark of viral infections^53^. J2 antibody staining, that specifically detects dsRNA, revealed that while some staining was present in the cytoplasm of uninfected cells, a distinct nuclear signal was exclusively present in all infected clones (Fig.3B). In order to ascertain that the J2 staining is a result of dsRNA originating from viral genomes, we implemented single molecule Fluorescence *In Situ* Hybridization (smFISH) and single molecule inexpensive FISH (smiFISH)^54^ to specifically detect each viral genome. J2 and FISH co-staining revealed a clear nuclear colocalization of both signals in all infected cells (Fig.3C). These data show that we can specifically stain viral genomes in infected cells and that the dsRNA signal is most likely originating from viral genomes.

**Figure 3:**
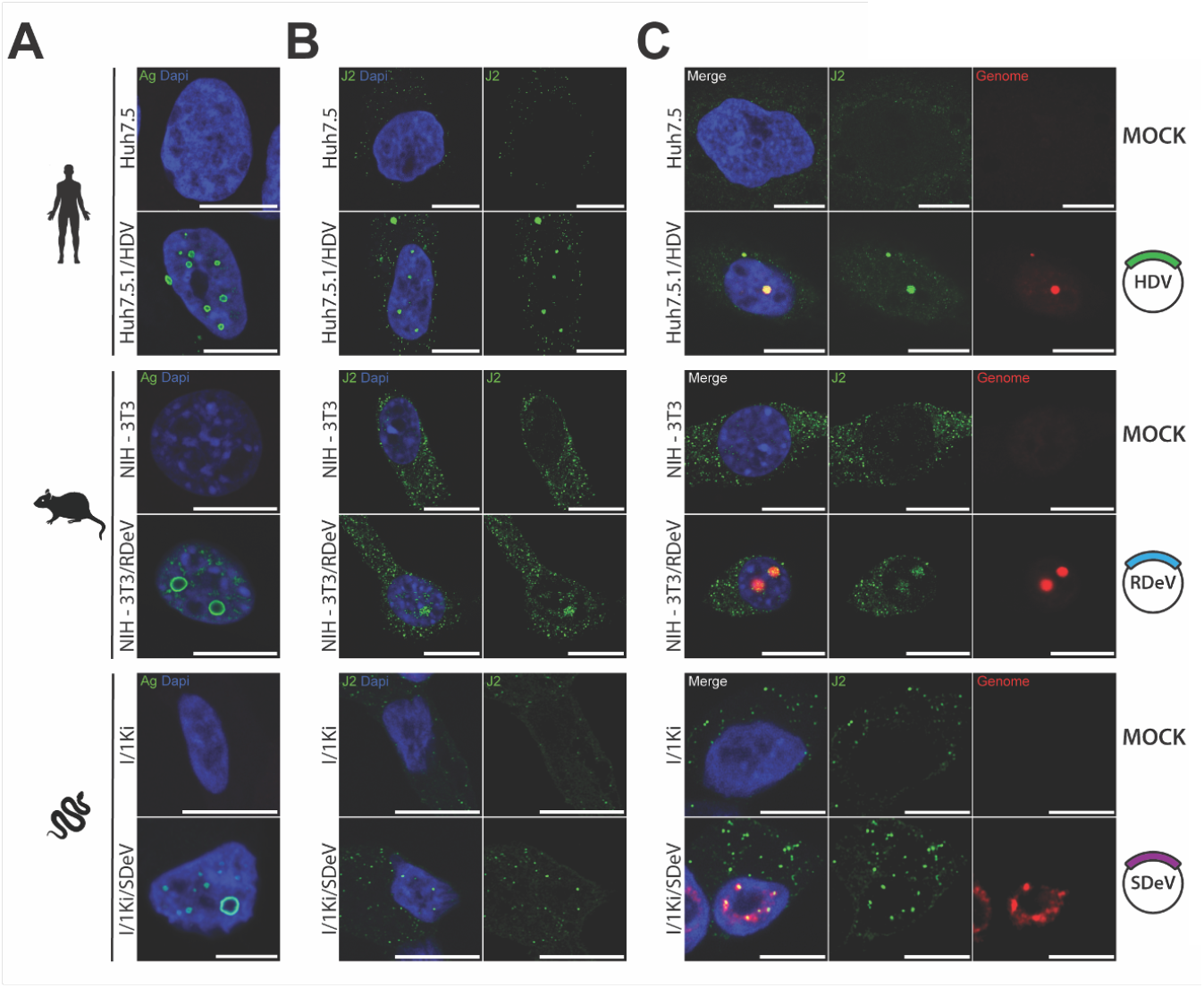
Kolmiovirid’s DAgs and genomes subcellular localization. Cell lines persistently replicating HDV, RDeV and SDeV were plated on microscopy slides and fixed to visualize DAg and genome subcellular localization in Huh7.5.1 replicating HDV, NIH-3T3 replicating RDeV and I/1Ki replicating SDeV. Non-infected cell lines served as negative controls. Nuclei (in blue) were stained using DAPI and representative confocal images are shown for A) Detection of the DAgs by immunofluorescence using the Ig-Patient1 serum (in green). B) Detection of double stranded RNA by immunofluorescence using the J2 antibody (in green). C) Co-detection of dsRNA by immunofluorescence using the J2 antibody (in green) and delta genomes by sm or smiFISH (in red). *Scale bars* 10 *μ*m for Huh7.5, Huh7.5.1/HDV, NIH-3T3 and NIH-3T3/RDeV and 5 *μ*m for I/1Ki and I/1Ki/SDeV. Cells were imaged on a LSM980 confocal microscope (Zeiss) and analyzed using ImageJ (version 2.9.0).

Interestingly, although DAg staining was almost exclusively nuclear for all three viruses, we noticed a heterogeneity in the distribution pattern of viral proteins in infected nuclei. We could classify the observed DAgs distributions in three distinct patterns (Fig.4A), resembling what has previously been observed with HDV^51,52^: 1-a “diffuse” localization throughout the nucleus, 2-a concentrated signal in foci distributed equivalently throughout all Z-stacks that we termed “dense hubs” and 3-concentrated signal in particular foci that form ring-like tructures, devoid of staining in focal planes positioned in the middle of the foci, that we termed “hollow hubs” (Fig.4A). Quantification of these patterns in HDV, RDeV and SDeV infected cells revealed that the “diffuse” pattern is the least abundant in most cell lines (Fig.4B), while viral protein hubs are the most frequent, with the “hollow hub” pattern represented in at least half of all counted cells for all three viruses (Fig.4B). The number of observed hubs per cell varied greatly between the three viruses (Fig.4C), with RDeV infected cells containing the least (mean of ∼4 hubs/cell) and SDeV infected cells containing the most (mean of ∼18 hubs/cell) (Fig.4C).

**Figure 4:**
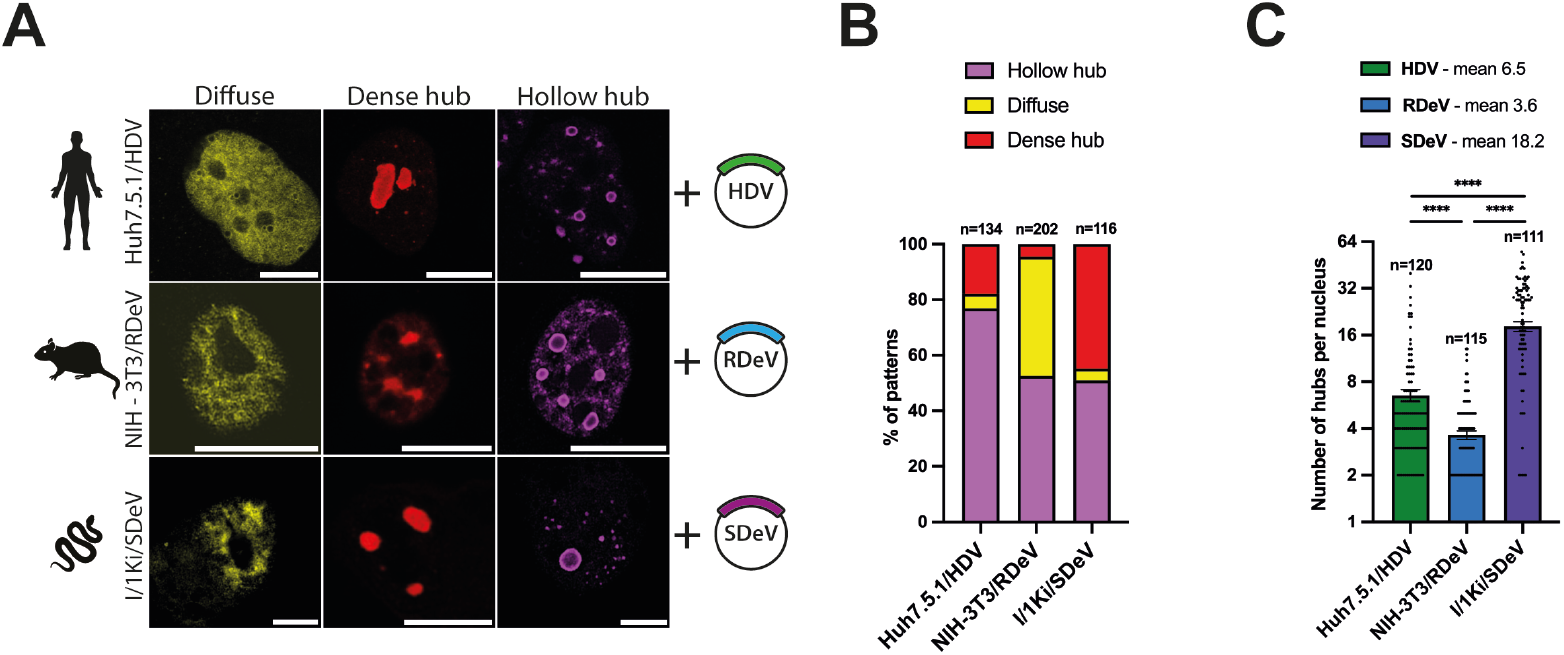
DAgs form different patterns in nuclei of infected cells. A) Mock or infected cells (Huh7.5.1 replicating HDV, NIH-3T3 replicating RDeV and I/1Ki replicating SDeV) were plated on microscopy slides and fixed to visualize DAgs structures in the nucleus. Representative confocal images are shown. DAgs (in yellow, red or magenta, depending on the pattern) were detected by immunofluorescence, *scale bars* 10 *μ*m for Huh7.5.1/HDV and NIH-3T3/RDeV and 5 *μ*m for I/1Ki/SDeV. Cells were imaged on a LSM980 confocal microscope (Zeiss) and analyzed using ImageJ (version 2.9.0). B) Quantification of the ratio between the three patterns described in panel A in Huh7.5.1/HDV, NIH-3T3/RDeV and I/1Ki/SDeV. Randomly selected fields of each cell line from at least 3 independent plating events were manually counted. C) Number of hubs (dense and hollow combined) in Huh7.5.1/HDV, NIH-3T3/RDeV and I/1Ki/SDeV. On the same randomly selected fields used for quantification in panel B, the number of hubs per cell was manually counted and plotted. Mean values and SEM (unpaired 2-side Student *t* test, with Welch’s correction, **** *P* < .0001).

Because the nuclear hollow hub is the most frequently observed pattern, we sought to determine the position of viral genomic RNA relative to DAg proteins in these structures using smFISH coupled to IF (smFISHIF) of infected cells. Results from these experiments showed a very similar 3D structural organization for all three viruses. In fact, the aforementioned “hollow hubs” turned out to be packed with viral RNA (Fig.5). Indeed, imaging of these hubs along the Z-axis shows a peculiar 3D organization, where viral RNA is concentrated in the middle of the hubs, surrounded by viral protein staining (Fig.5). To confirm this organization, we quantified protein and RNA signals along the X-axis from images at different Z-stacks of representative “hollow hubs”. Two overlapping signals were detected at the apical and basal poles (Fig.5) whereas the middle of the hub shows a peak in RNA signal intensity surrounded by two antigen signal peaks (Fig.5). 3D reconstitutions of these hubs, show spherical structures full of viral RNA surrounded by a “shell” of viral proteins (Sup Video 1-3).

**Figure 5:**
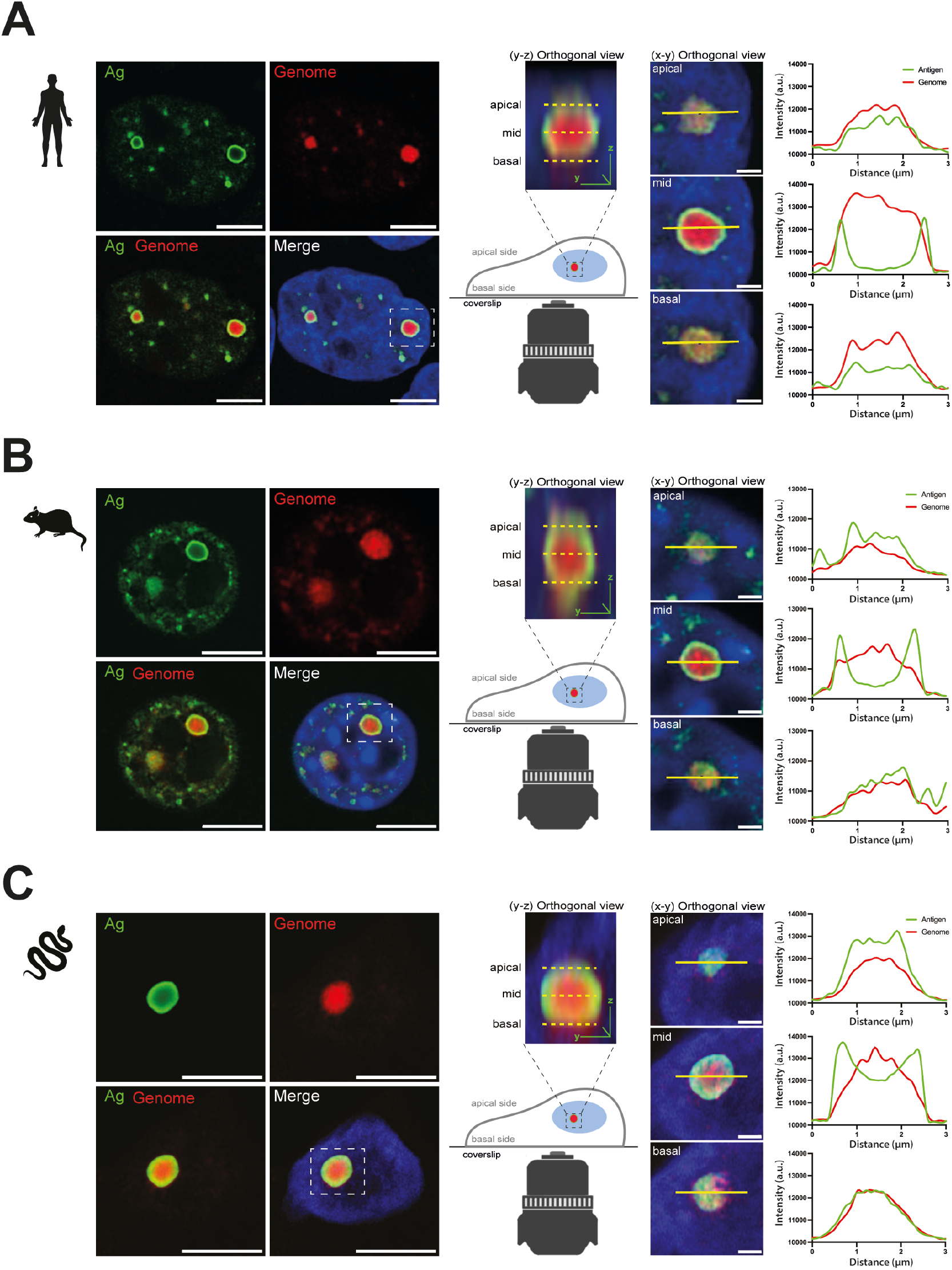
Viral hubs in infected cells. Mock or infected Huh7.5.1 replicating HDV (A), NIH-3T3 replicating RDeV (B) and I/1Ki replicating SDeV (C) were plated on microscopy slides and fixed to visualize DAgs and RNA localization. Representative confocal images are shown. Orthogonal view (Y,Z) showing the organization of the hubs along the Z-axis are shown and used to define the basal, mid and apical planes of the presented hubs. Fluorescence intensity of the antigen and genome signals were determined along the X-axis at the basal, mid and apical planes of the hubs and plotted adjacent to the corresponding orthogonal views (X, Y). Nuclei (in blue) were stained using DAPI, DAgs (in green) were detected by IF staining and delta genomes (in red) were detected by sm or smiFISH, *scale bars* 10 *μ*m for Huh7.5.1/HDV and NIH-3T3/RDeV confocal panels, 5 *μ*m for I/1Ki/SDeV confocal panels and 1 *μ*m for all X, Y orthogonal views. Cells were imaged on a LSM980 confocal microscope (Zeiss) and analyzed using ImageJ (version 2.9.0).

Host RNA Polymerase II (RNAPII) is thought to be the main polymerase responsible for HDV RNA amplification^25,55^. Thus, we investigated whether observed viral RNA-protein structures correspond to transcription foci enriched in RNAPII. RNAPII staining was distributed throughout nuclei as expected (Fig.S2). However, even though viral protein and RNA Pol II staining co-localized in some areas, no clear enrichment of RNAPII signal was observed in viral protein hubs (Fig.S2), similar to what has been observed earlier for HDV^50^.

### HDV, RDeV and SDeV tail vein injections and viral replication in vivo

The development of a mouse model to study *Kolmioviridae* replication and associated pathogenesis would have obvious benefits. Chang *et al*.^56^ were able to initiate HDV replication in mouse hepatocytes *in vivo* by hydrodynamic tail vein injection (HDTV) of plasmid DNA harboring HDV genome^56^ (Fig.6A). We applied this method to verify if RDeV and SDeV, similarly to HDV, can replicate in hepatocytes *in vivo*.

**Figure 6:**
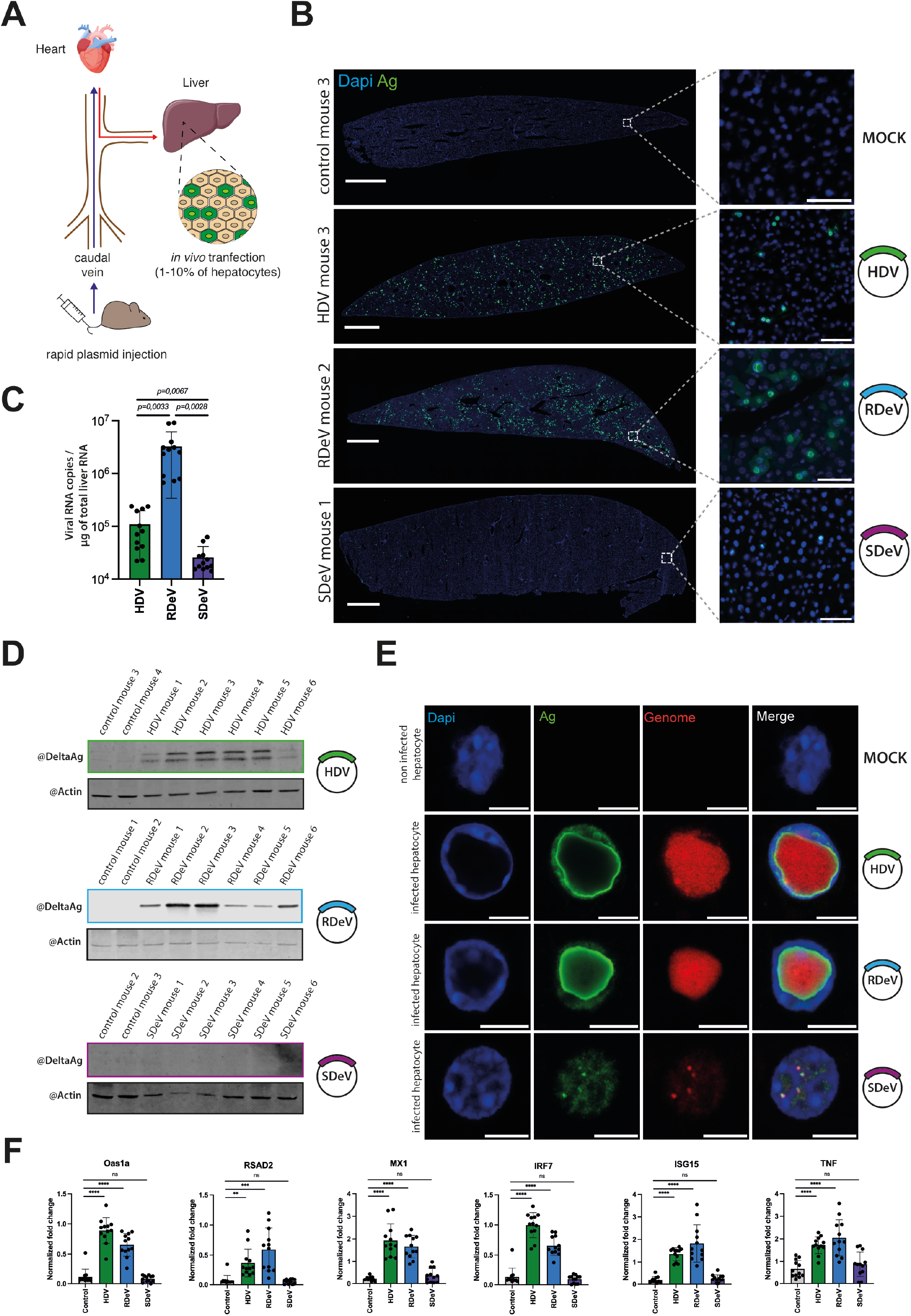
Deltaviruses’ replication *in vivo*. Mice injected with empty pCDNA3.1 or pCDNA3.1 encoding dimers of the HDV, SDeV or RDeV genomes were euthanized and dissected 9 days post-injection. A) Schematic representation of hydrodynamic tail vein injections in mice. B) Immunofluorescence detection of the DAgs in mice liver sections. Liver samples were collected, sliced (10 mm) and fixed to visualize DAg expression in hepatocytes. Representative images are shown. Nuclei (in blue) were stained using DAPI and DAgs (in green) were stained by IF, *scale bar* 1 mm. The boxed areas are shown magnified, *scale bar* 50 *μ*m. Slices were imaged on an Axioscan 7 (Zeiss) and analyzed using Zen Blue (Zeiss, version 3.7) and ImageJ (version 2.9.0). C) qPCR analysis of viral RNA copy numbers in dissected mice liver. Liver samples were collected and total extracted RNAs were used to synthetize cDNAs for qPCR analysis of the viral genomes. RNA levels were normalized using the level of the 18S ribosomal RNAs for each sample and viral RNA copy numbers were calculated using a serial viral plasmid dilution. Mean values and SD were calculated for 6 independent injections (i.e., 6 mice) with technical duplicates (n = 6 with technical duplicates for each, unpaired 2-side Student *t* test, with Welch’s correction, **** *P* < .0001). D) Western blot analysis of the DAgs’ expression in dissected mice livers. Liver samples were collected and protein extracts were analyzed by western blot for DAg and β-actin expression. E) smFISHIF detection of the DAgs and viral genomes in mice liver. Liver samples were collected, sliced (10 mm) and fixed to visualize DAg and genome localization in hepatocytes. Representative images are shown. Nuclei (in blue) were stained using DAPI, DAgs (in green) were detected by IF staining and delta genomes (in red) were detected by sm or smiFISH, *scale bar* 10 *μ*m. Slices were imaged on a LSM980 confocal microscope (Zeiss). F) qPCR analysis of ISG fold change expression in dissected mice liver. Liver samples were collected and total RNAs extracted and used to synthetize cDNAs for qPCR analysis of a selection of ISGs. RNA levels were normalized using the level of the 18S ribosomal RNAs for each sample. Mean values and SD were calculated for 6 independent injections (i.e., 6 mice) with technical duplicates (n = 6 with technical duplicates for each, unpaired 2-side Student *t* test, with Welch’s correction, ** *P* < .01, *** *P* < .001, **** *P* < .0001, ns: not statistically significant.).

We performed HDTV on 24 C57BL/6J female mice in total, with plasmids harboring either HDV (n=6), RDeV (n=6) or SDeV (n=6) genomes, or empty pCDNA3.1 as mock control (n=6) (Fig.6A). We sacrificed the animals 9 days post-injection and collected liver samples. IF staining of DAgs on liver sections showed no signal in mock injected mice, a very weak signal in SDeV injected animals and a clear nuclear signal in hepatocytes of HDV and RDeV injected mice (Fig.6B). These results are in agreement with viral RNA quantification determined by RT-qPCR (Fig.6C). Indeed, while RDeV RNA was the most abundant, HDV RNA was ∼1.5 log lower and SDeV RNA was 2 logs less abundant than RDeV RNA (Fig.6C). Western blot analysis detected both small and large HDAgs as well as RDAgs in all injected HDV and RDeV mice, however SDAgs was undetectable in animals injected with the SDeV plasmid (Fig.6D). Altogether, these results show that similarly to HDV^56^, RDeV can efficiently replicate in mouse hepatocytes, reaching even higher levels than HDV (Fig.6C). SDeV on the other hand, replicates very poorly in mouse liver (Fig.6B-D), consistent with our *in vitro* data (Fig.1D).

Strikingly, our IF results on stained liver slices of RDeV and HDV injected mice showed massive hubs with unusual nuclear DAg staining (Fig.6B). Using smFISHIF and confocal microscopy to take a closer look at these hubs, we observed that both HDV and RDeV positive nuclei accumulated large amounts of viral RNAs, forming massive hubs of RNA surrounded by DAg staining (Fig.6E), reminiscent in their 3D organization to the hubs we observed in culture models (Fig.5) albeit much larger in size (Fig.6E, Sup Video 4-6). Remarkably, in the majority of HDV and RDeV positive cells, RNP hubs occupied up to ∼75% of the host cell’s nuclear volume, displacing the DNA counterstain by DAPI to nuclear borders (Fig.6E, Sup Video 4-6). Interestingly, in the very few SDeV positive cells that we were able to find, the smFISHIF staining revealed much smaller RNP foci (Fig.6E, Sup Video 6).

### Unraveling human, rodent and snake Kolmioviridae replication to anticipate cross-species transmission

Finally, we wished to verify if *Kolmioviridae* replication via HDTV could recapitulate immune gene induction observed with HDV^17^ in several infection models^57–60^. RT-qPCR probing a selection of interferonstimul ated genes (ISGs) and tumor necrosis factor alpha (TNFα) mRNA was performed on RNA extracted from livers from different mouse groups (Fig.6F). Mx1, Oas1a, RSAD2, ISG15, IRF7 and TNFα mRNAs were all upregulated upon HDV and RDeV injection, however, no induction was observed after SDeV injection, suggesting that the immune gene induction is likely the result of kolmiovirid replication.

## Discussion

The discovery of HDV-*like* sequences in duck cloacal samples and in various boa constrictor organs, prompted several laboratories to examine and identify similar sequences across a wide range of taxa^6–12,15^. The findings led to the creation of the novel realm *Ribozyviria* with the family *Kolmioviridae* to host the genus *Deltavirus* and seven other genera^16^. As only a minority of the studies on novel kolmiovirids made efforts towards molecular characterization of their replication^9,11,14^, our understanding of the kolmiovirid biology relies on the seminal research performed on HDV. Importantly, the cross-species transmission potential of kolmiovirids^10^ coupled to their ability to form infectious particles with envelope proteins of a variety of helper viruses ^13,32^ raises an important question: can these minimal RNA viruses replicate in any animal cell they access? Here, by comparing molecular replication hallmarks of two recently identified kolmiovirids, RDeV and SDeV, to HDV, we reveal differential and convergent features governing their interaction with their host, and provide estimations of their host range, and thus host-shifting potential^10^.

To allow unbiased comparison, we constructed infectious clones for HDV, RDeV, and SDeV in the same orientation and expression vectors and studied their replication in nine different animal cell lines. We show that while RDeV replicated in all tested cell lines, HDV replicated in the majority of the mammalian cell lines and SDeV replicated most efficiently in snake cells (Fig.1C-G). This indicates that while RDeV appears to be a generalist, host specific factors control SDeV and HDV replication. In the future, determining sequences in RDeV genome (e.g. antigen, promotor sequences, ribozymes) conferring this virus its “super-replicator” status would shed light on important aspects of kolmiovirid replication.

ADAR-1-dependency of HDV but neither SDeV nor RDeV, illustrates the requirement for host specific factors (Fig.2). While SDeV produces a single form of SDAg in cultured snake cells (Fig.1E) and in infected snakes^13,14^, we observed two forms of the SDAg in human Huh7.5 and HEK293T cells (Fig.1C). Using ADAR-1-KO or overexpressing cell lines, as well as a SDeV reporter infectious clone, we showed that the two SDAg species are not the result of an editing event in the SDeV genome (Fig.2). Because neither SDeV nor RDeV appear to require ADAR-1 (Fig.2), it is tempting to speculate that co-evolution with HBV, particularly HBsAg, made ADAR-1 editing useful for HDV that co-opted its function. The ADAR-1 editing allows for the prenylation of cysteine 211 on the L-HDAg, that anchors it to cellular membranes, facilitating its interaction with HBsAg and the assembly of HDV infectious particles^30,61–63^. Our results suggest that, in addition to ADAR-1 editing, divergent host factor dependencies and specific viral sequences likely contribute to kolmiovirid’s host-specificity. Our data call for more systematic studies addressing host-shifting potential of kolmiovirids assessing which virus would potentially be more prone to zoonoses.

The analysis of kolmiovirid replication in persistently infected cells through IF revealed three DAg staining patterns similar for all viruses. Simultaneous viral RNA and DAg detection of the major hollow hub pattern through smFISHIF followed by high-resolution microscopy, revealed that the observed hubs are in fact full of a viral RNA. Infected cells showed a peculiar RNA-protein structure, formed by RNA condensates surrounded by a layer of DAg. This organization seems to be a defining feature of kolmiovirid accumulation in infected nuclei, as it was also observed *in vivo* in infected mice liver (Fig.6E). Interestingly, RNA-protein hubs observed in mouse hepatocytes were often massive, occupying the majority of infected nuclei. One hypothesis explaining the difference between the observed patterns and the size of different hubs relates to cell cycle progression. Specifically, the hubs would be smaller in size and in higher number in immortalized cell cultures, because of their dilution after each cell division. In contrast, continuous viral replication in non-dividing mouse hepatocytes would cause these hubs to grow and presumably coalesce, reaching very large sizes. Admittingly, the function of these hubs during the viral lifecycle is unclear but one could speculate that they can either be viral assembly sites or active viral replication factories. Costaining for viral Ag and RNAPII did not reveal an enrichment of RNAPII in these structures, however we used an antibody recognizing hypo-phosphorylated RNAPII, so one cannot rule out that the virus might use a hyper-phosphorylated RNAPII form for its RNA-dependent RNA polymerase function^55,64^. Future experiments with labeled UTP will indicate if RNA transcription is active in these structures. Curiously, the organization of these structures is reminiscent of liquid-liquid phase separations (LLPS) observed in membraneless organelles, such as the nucleolus^65^ or several viral inclusion bodies (IB)^66,67^. Indeed, many viruses, including negative-strand RNA viruses^68^, such as Rabies virus (RABV)^69^, Vesicular stomatitis virus (VSV)^70^, Measles virus (MeV)^71^ and influenza A virus (IAV)^72^ are known to form LLPS in the cytoplasm of infected cells to promote viral replication or genome assembly^73^. Interestingly, a common feature of these virus-induced LLPS is that their formation requires, apart from an RNA component, proteins containing RNA binding motifs, oligomerization domains and domains with intrinsically disordered regions (IDRs)^73^, which are all present in DAgs (Fig.1A). In the future, tracking the morphology of the observed kolmiovirid hubs by live microscopy approaches (e.g. ability to fuse, fluorescence recovery after photobleaching), would determine if the observed hubs have liquid- or gel-like properties^74^.

Finally, our HDTV experiments show that unlike RDeV and HDV, SDeV is not able to efficiently replicate in mouse hepatocytes *in vivo*, in agreement to what is observed in cell culture. This suggests that the observed differences likely reflect the ability of RDeV and HDV, but not SDeV, to initiate replication in mice, which would subsequently dictate the induction of ISGs^57,60,75^(Fig.6). In conclusion, our results suggest differences in the cross-species transmission potential between kolmiovirids. Importantly, amongst viruses evaluated in our study, RDeV appears to be omnipotent in initiating replication in cells of various animal species.

## Supporting information

Supplementary video 1: 3D reconstitution of an infected Huh7.5.1/HDV nucleus. Confocal Z-stack images from the infected nucleus presented in Figure 5A

Supplementary video 2: 3D reconstitution of an infected NIH-3T3/RDeV nucleus. Confocal Z-stacks images from the infected nucleus presented in Figure 5

Supplementary video 3: 3D reconstitution of an infected I/1Ki/SDeV nucleus. Confocal Z-stacks images from the infected nucleus presented in Figure 5C

Supplementary video 4: 3D reconstitution of an HDV infected mouse hepatocyte nucleus. Confocal Z-stacks images from the infected nucleus presented in

Supplementary video 5: 3D reconstitution of an RDeV infected mouse hepatocyte nucleus. Confocal Z-stacks images from the infected nucleus presented in

Supplementary video 6: 3D reconstitution of an SDeV infected mouse hepatocyte nucleus. Confocal Z-stacks images from the infected nucleus presented in

## Acknowledgements

We would like to thank the following: Montpellier Bio-Campus platforms for contributing to this project: Montpellier Resources Imagerie (MRI) for microscopy and flow-cytometry, Réseau d’Histologie Experimentale de Montpellier (RHEM) for histology, Réseau des Animaleries de Montpellier (RAM) for animal care. Some experiments were performed in CEMIPAI (UAR 3725 CNRS/Montpellier University) BSL3 Facility. We would like to thank Troels Scheel, Jean-Luc Battini, Virginia Pimmett, Jean-Pierre Vartanian, Caroline Goujon, Florence Rage and Anja Kipar for providing expertise and reagents for the study. We would like to thank Dominique Helmlinger, Fabrice Caudron, Joe McKellar and Urszula Hibner for critical reading of the manuscript. The project was funded by the ATIP-AVENIR program (CNRS-INSERM), the MUSE (Montpellier Université d’Excellence) grant, by the Deutsche Forschungsgemeinschaft (DFG, German Research Foundation) to D.G. project number 197785619 (SFB 1021) and the European Research Council (ERC) starting grant 101039538 – DELV.

## Materials and Methods

### Sequence analyses and computational protein structure prediction

The multiple protein sequence alignment shown in Fig.1A was computed by Clustal Omega on the EMBL-EBI server. Computational meta-disorder predictions and consensus secondary structure prediction were obtained from the Dismeta webserver^76^. Conservation scores based on alignment of sequences from HDAg (genotype 1 – HDV isolate Taylor), Tome’s spiny rat virus 1 (TSRV-1) delta antigen from isolate 0180 (rodent DAg) and Swiss snake colony virus 1 (SwSCV-1) delta antigen from isolate F18-5 (snake DAg) were calculated using AL2CO^77^ with a 10-residue sliding average, as implemented in Chimera^78^. Structure predictions of HDAg, RDAg and SDAg were performed using a SBGrid consortium installation of AlphaFold multimer version 2.3 running on a local server equipped with a NVIDIA Tesla A100 GPU^79,80^. The full databases were used, with max_template_date = 2022-12-22. All other parameters were left to their default values.

### Plasmids and Cloning

pcDNA3.1 plasmids encoding dimers of the human (Genebank accession number M21012.1) and rodent (Genebank accession number : MK598004) deltaviruses’ genomes where previously described^9^. pcDNA3.1 plasmid encoding a dimer of snake deltavirus (Genebank accession number MH988742) was generated by PCR amplification of the snake deltavirus dimer insert from pCAGGS-2xSDeV-fwd^13^ and ligating it in a EcoRV/XbaI linearized pCDNA3.1. Clones were sequenced and plasmid in the forward orientation were selected and amplified for further use. PlentiCMVPuro plasmids (Addgene, #17452) encoding the FLAG tagged human, rodent, avian, toad and snake delta (SDAg-S and SDAg-L) viruses’ antigens were generated by PCR amplification of the antigens coding sequences with primers listed in Table 1, from pCAGGS 1.2x plasmids of HDV, RDeV, SDeV, avian and toad deltaviruses described in Szirovicza *et al*.^14^. The amplified fragments were then cloned into PlentiCMVPuro (Addgene, #17452) plasmids by Gibson assembly from NEB (#E2611L). The SDeV infectious clones with HA-tag C-terminally to either S-SDAg or the putative L-SDAg were constructed as 1.2x genome inserts in genomic orientation into pCAGGS vector similarly to what has been described for 1.2x genome SDeV^14^. The ADAR-1 p110 and ADAR-1 p150 coding sequences were amplified by PCR from pcDNA3.1 cDNA plasmids kindly provided by J.P. Vartanian and cloned into the pLentiCMVPuro DEST vector (Addgene, #17452) using Gibson assembly. All primers used for cloning are listed in Supplementary Table 1.

### Cell culture

Huh7.5 (Cancerous Human hepatocytes, RRID:CVCL_7927), HEK293T (transformed human embryonic kidney cells RRID:CVCL_0063), A549, NIH-3T3 (spontaneously immortalized NIH Swiss mouse embryonic fibroblasts, RRID:CVCL_0594), MCA-RH 7777 (cancerous buffalo rat hepatocytes, RRID:CVCL_0444), FEA (spontaneously immortalized cat embryonic fibroblasts, RRID:CVCL_UG17), CRFK (spontaneously immortalized cat kidney epithelial cells, RRID:CVCL_2426), and Vero (spontaneously immortalized green monkey kidney epithelium, RRID:CVCL_0059) cells were maintained in Dulbecco’s modified Eagle’s medium (DMEM; Gibco #41965-039) supplemented with 10% heat-inactivated fetal calf serum (Sigma-Aldrich #F75424), and 100 mg/mL of normocin (InvivoGen, #ant-nr-2) and grown in cell culture flasks at 37 °C in a humidified incubator containing 5% CO2. I/1Ki (*boa constrictor* kidney cells)^81^ and I/1Ki-Δ (refered to as I/1Ki/SDeV in this paper)^13^ were maintained in Minimal Essential Medium Eagle (MEM; Gibco #31095-029) supplemented with 10% heat-inactivated fetal calf serum (Sigma-Aldrich #F75424), and 100 mg/mL of normocin (InvivoGen, #ant-nr-2), and grown in cell culture flasks at 30 °C in a humidified incubator containing 5% CO2. All cell lines used in the study were mycoplasma-free and were checked regularly for mycoplasma contamination.

### Transfections

Huh7.5, HEK293T, A549, NIH-3T3, MCA-RH 7777, FEA, CRFK, and Vero cells were transfected with Jet Pei reagent (Polyplus; #101000020) using 1 μg of plasmid DNA and according to the manufacturer’s instructions. I/1Ki cells were transfected with Lipofectamine 3000 (Invitrogen, #L3000008) using the reverse transfection method. 1.5 μL Lipofectamine 3000 was diluted in 25 μL of Opti-MEM Medium, and 1 μg of plasmid DNA was diluted with 2 μL P3000 reagent in 25 μL of Opti-MEM. Both solutions were mixed, incubated for 15min at room temperature (RT) and added to 1.5×10^5^ cells suspended in 1mL of culture medium. The cells were kept in suspension with the transfection mix for 15 to 30 minutes RT before being seeded in 24 well plates. Transfection medium was replaced for culture medium 6h post-seeding.

### Establishment of ADAR-1 KO cell lines

To generate ADAR-1 p110 and p150 KO cell lines, the following primers corresponding to guide RNAs targeting early exons of ADAR-1 were cloned into the *Sp*Cas9-expressing lentiviral vector lentiCRISPRv2^47^: ADAR-1 p110 and p150 Fwd: 5’-CACCGAATTGACATGGAAAGGCAGG-3’; and Rev: 5’-AAACCCTGCCTTTCCATGTCAATTC-3’; and ADAR-1 p150 Fwd: 5’ CACCGAATTGACATGGAAAGGCAGG-3’; and Rev: 5’-AAACCCTGCCTTTCCATGTCAATTC-3’. Lentiviral particles pseudotyped with the VSV-G protein were produced by co-transfecting HEK293T cells in 6 well plates with 3 mg lentiCRISPRv2 vector^47^, 1.5 mg VSV-G Env expression vector pMD2.G (Addgene #12259) and 1.5 mg Gag-Pol expression vector psPAX2 (Addgene #12260) vector using a standard calcium chloride transfection protocol. Viral supernatants were harvested 48 h after transfection, filtered (0.45 μm), and stored at −80°C or used directly for transduction. Huh7.5 and HEK293T transduced cell lines were selected in 1 μg/mL or 3.5 μg/mL of puromycin respectively for at least 5 days prior to use in assays. Thereafter, protein lysates were collected from the transduced cells and protein levels of the ADAR-1 p110 and p150 were assessed by immunoblotting.

### Establishment of ADAR-1 overexpressing cell lines

To generate ADAR-1 p110 and p150 overexpressing cell lines, lentiviral particles pseudotyped with the VSV-G protein were produced by cotransfecting HEK293T cells in 10cm dishes with 5 mg pLentiCMVPuroDEST vector (Addgene, #17452), 2 mg VSV-G Env expression vector pMD2.G (Addgene #12259) and 2 mg Gag-Pol expression vector psPAX2 (Addgene #12260) using Jet Pei reagent (Polyplus; #101000020) according to the manufacturer’s instructions. Viral supernatants were harvested 48 h after transfection, filtered (0.45 μm), and stored at −80°C or used directly for transduction. Huh7.5 and HEK293T transduced cell lines were selected in 1 μg/mL or 3.5 μg/mL of puromycin respectively for at least 5 days prior to use in assays. Thereafter, protein lysates were collected from the transduced cells and protein levels of the ADAR-1 p110 and p150 were assessed by immunoblotting.

### Establishment of HDV and RDeV expressing cell lines

To establish an Huh7.5.1 cell line persistently replicating HDV and a NIH-3T3 cell line persistently replicating RDeV both cell lines were cotransfected with an mCitrine encoding plasmid kindly provided by Eric Kremer’s lab and a pcDNA3.1 plasmid encoding a dimer kolmiovirid coding sequence in a 1:4 ratio, HDV genotype 1 (Taylor isolate -GenBank accession number M21012.1) and RDeV isolate 0183 (GenBank accession number MK598004) respectively. Two days post-transfection single clones were sorted into 96-well plates containing conditioned medium by Fluorescence Activated Cell sorting (FACS). Cell clones were amplified, and screened for the presence of HDV or RDeV genomes and antigens by RT-qPCR and western blot. Clones were selected for this study and grown in the same conditions as their respective parental cell lines.

### Animal models

All reported animal procedures were carried out in accordance with the rules of the French Institutional Animal Care and Use Committee and European Community Council (2010/63/EU). Animal studies were approved by institutional ethical committee (Comité d’éthique en expérimentation animale Languedoc-Roussillon (#36)) and by the Ministère de l’Enseignement Supérieur, de la Recherche et de l’Innovation (Apafis #40209-2023010417589371 v3). Hydrodynamic tail vein injections were performed in 6 to 8 week-old C57Bl/6J female mice (Janvier), as previously described^82^. Briefly, 0.1 mL/g body weight of a solution of sterile saline containing plasmids of interest were injected into lateral tail vein over 8-10s. HDV / SDeV / RDeV, or empty pCDNA3.1 as control (12.5 μg) were injected together with pSBbi-RN (Addgene #60519) integrating reporter plasmid encoding dTomato (12.5 μg) and sleeping beauty transposase SB100X (Addgene #34879) (2.5 μg). pSBbi-RN (Addgene #60519) and pCMV(CAT)T7-SB100 (Addgene #34879) were gifts from Eric Kowartz and Zsuzsanna Izsvak, respectively. At 8-9 days post-injection, mice were fasted for 4-6 hours then sacrificed by anesthetic overdose with isoflurane. Livers were collected and cryopreserved in tissue freezing medium (TFM-5, Microm microtech) in liquid Nitrogen cooled isopentane following classical procedure. RNA and proteins were extracted from snap frozen samples of the liver caudate lobe.

### Antibody isolation from patient serum

Antibody targeting Hepatitis delta antigen (HDAg) were purified from serum of a cohort of HBV/HDV coinfected patients (n=6)^83^ using the MabTrap® Kit (Cytvia, #17112801) according to the manufacturer’s instructions. The antibody used for all subsequent DAgs stainings and western blot experiments, Ig-Patient1, was described earlier^40,41^. Human serum from patients with chronic HBV/HDV infection followed at the Strasbourg University Hospitals, Strasbourg, France, was obtained with informed consent.

### Western blot

Cells were washed with PBS and lysed with 1X RIPA buffer (Merck Millipore #20-188) supplemented with a protease inhibitor cocktail (Thermo Scientific #87785). Total protein samples were denatured in 1X Laemmli buffer (Bio-Rad #1610747) supplemented with 10% β-mercaptoethanol (Bio-Rad #1610710) for 5 min at 95°C, and loaded on 4– 20% Mini-PROTEAN® TGX gels (Bio-Rad #4561093). Electrophoresis was performed at 120V for 1 hour. Proteins were subsequently transferred onto PVDF membranes (Bio-Rad #10026933) for 7 min at 2.5A and 12V using a Trans-Blot® Turbo™ Transfer System (Bio-Rad #1704150EDU). Membranes were saturated in PBS (137 mM NaCl, 2.7 mM KCl, 10 mM Na_2_HPO_4_, 1.8 mM KH_2_PO_4_) 0.1% Tween 20 (Bio-Rad #1610781) (PBST) containing 5% dry milk (Régilait #731142) for 30 min at RT. Specific primary antibodies were incubated overnight at 4°C in PBST containing 2% dehydrated milk. Membranes were washed 3 times in PBST at RT and iRDye labeled specific secondary antibodies were incubated for 1 hour at RT in PBST in the dark. Membranes were washed 3 times in PBST at RT and iRDye labeled specific secondary antibodies were detected using the Odyssey® M Infrared Imaging System (LI-COR Biosciences #3350). Proteins from liver tissues were extracted from freshly dissected mouse liver tissue fragments, collected in 2 mL tubes, flash frozen in liquid nitrogen and treated with a lysis buffer (150mM NaCl, 50mM Tris pH7.5, 1% Triton X-100 (Bio-Rad #1610407), 1% SDS) supplemented with a protease and phosphatase inhibitor cocktail (Thermo Scientific, # 78430). The protein concentration of each lysate was measured using the PierceTM BCA Protein Assay Kit (Thermo Scientific #23227) and 20 mg of protein from each sample were denatured and treated as described above.

Mouse anti β-actin monoclonal antibodies (Invitrogen, MA5-11869) diluted 1:2,000 were used to detect β-actin protein as a loading control. Polyclonal antibodies extracted from patient sera (Ig-Patient1 for all blots except HDV and SDeV antigens detected in mice samples which were detected with Ig-Patient5) were used to detect the HDAg and RDAg. Rabbit serum/antiserum immunized with recombinant SDAg^7^ diluted 1:2,000 were used to detect SDAg. Rabbit anti-ADAR-1 monoclonal antibodies (Cell Signaling Technology D7E2M #14175) and polyclonal antibodies (ThermoFisher Scientific #A303-883A-T) diluted 1:2,000 were simultaneously used to detect ADAR-1 p110 and p150. iRDye 800CW-conjugated goat anti human IgG secondary antibodies (LI-COR Biosciences #926-32232), iRDye 680RD-conjugated donkey anti-mouse IgG secondary antibodies (LI-COR Biosciences #926-68072) and iRDye 800CW-conjugated donkey anti-rabbit IgG secondary antibodies (LI-COR Biosciences #926-32213) diluted 1:10,000 were used to detect human sera, mouse β-actin antibodies, rabbit ADAR-1 antibodies and rabbit SDAg anti-serum, respectively.

### Immunofluorescence staining applied to cells

Cells were grown on microscope cover glasses (Marienfeld #0102052) in 6-well plates, washed 3 times with PBS (Eurobio #CS1PBS01-01) then fixed and permeabilized using a 4% paraformaldehyde (Electron Microscopy Sciences # 15714), 0.2% Triton X-100 (Bio-Rad #1610407) PBS solution for 20 min at RT. Cover glasses were washed 3 times with a 0.1 M Tris-HCl, 0.15M NaCl solution and saturated in saturation buffer: PBS, 0.1% Triton X-100, 2% BSA (Sigma-Aldrich #A3059) solution for 30 min at RT before overnight incubation with primary antibodies in the same buffer at 4°C. Cover glasses were subsequently washed 3 times with a 0.1M Tris-HCl, 0.15M NaCl solution and incubated in the dark with secondary antibodies for 2 hours in the saturation buffer at RT and in the dark. Slides were incubated with 300nM DAPI (Invitrogen #D21490) in a 0.1M Tris-HCl, 0.15M NaCl solution for 15 min at RT and in the dark and washed 3 times 5 min in a 0.1M Tris-HCl, 0.15M NaCl solution. Cover glasses where then mounted in ProLong Gold antifade reagent (Invitrogen #P36930), left to polymerize overnight at RT and in the dark then sealed with nail polish.

Polyclonal antibodies extracted from patient sera (Ig-Patient1) diluted 1:500 were used to detect the various delta antigens, monoclonal mouse anti-dsRNA J2 IgG (Jenna Bioscience, # RNT-SCI-10010200) diluted 1:500 was used to detect dsRNA and monoclonal mouse anti-polR2A IgG (8WG16, Invitrogen, #MA1-26249) diluted 1:50 was used to detect RNAPII. Alexa Fluor 488-conjugated goat anti-human IgG secondary antibody (Invitrogen, #A11013) diluted 1:500 was used to detect human primary antibodies, Alexa Fluor 488-conjugated goat anti-mouse IgG secondary antibody (Invitrogen, #A11001) and Alexa Fluor 555-conjugated donkey anti-mouse IgG secondary antibody (Invitrogen, #A31570) diluted 1:500 were used to detect mouse primary antibodies.

### Single molecule in situ hybridization immuno-fluorescence (smFISHIF) staining applied to cells

Cells were grown on microscope cover glasses (Marienfeld #0102052) in 6 well-plates, washed 3 times with PBS (Eurobio #CS1PBS01-01) and fixed and permeabilized using a 4% paraformaldehyde (Electron Microscopy Sciences # 15714), 0.2% Triton X-100 (Bio-Rad #1610407) PBS solution for 20 min at RT. Cover glasses were washed 3 times in PBS, saturated in a PBS 0.5% ultrapure BSA (ThermoFisher Scientific #AM2616) solution for 30 min at RT and washed for 20min in a 10% formamide (Merck, #F9037), 2X SSC (Invitrogen, #AM9770) solution before overnight incubation in a 10% formamide, 2X SSC, 8% dextran sulfate (Sigma-Aldrich, #D8906), 0.34mg/mL *E. coli* tRNA (Roche, #10109541001), 1mM vanadyl ribonucleoside complex (VRC, merck-millipore #R3380), 0.01% ultrapure BSA buffer containing 125nM Cy5 coupled smFISH probe mix or 2% smiFISH Cy5 duplexed probe mix (probes used are listed in Supplementary table 2) and the required antibodies at 37°C and in the dark. FLAP-Y containing smiFISH probes were previously duplexed with Cy5 conjugated FLAP-Y probes in the following conditions: 68ng/uL probe mix, 7 mM Cy5 conjugated FLAP-Y probes, 10% NEBuffer 3 (New England Biolabs, #B7003S) in a thermocycler at 85°C for 3 min, 65°C for 3 min and 25°C for 5 min. Cover glasses were subsequently washed twice for 15 min at 37°C in a 10% formamide, 2X SSC solution, incubated for 1h in a 0.01% ultrapure BSA, 10% formamide, 2X SSC containing the required secondary antibodies and then in a 300nM DAPI (Invitrogen #D21490), 10% formamide, 2X SSC solution, for 15 min at 37°C in the dark and washed 3 times 5 min in a 2X SSC, 0.1% Tween 20 (Thermo Scientific, #J20605-AP), PBS solution. Cover glasses were then mounted in ProLong Gold antifade reagent (Invitrogen #P36930), left to polymerize overnight at RT and in the dark then sealed with nail polish.

Polyclonal antibodies extracted from patient sera (Ig-Patient1) diluted 1:500 were used to detect the various delta antigens, monoclonal mouse anti-dsRNA J2 IgG (Jenna Bioscience, # RNT-SCI-10010200) diluted 1:500 was used to detect dsRNA. Alexa Fluor 488-conjugated goat anti-human IgG secondary antibody (Invitrogen, #A11001) diluted 1:500 was used to detect human primary antibodies, Alexa Fluor 488-conjugated goat antimouse IgG secondary antibody (Invitrogen, #A11001) was used to detect mouse primary antibodies.

### Immunofluorescence staining applied to tissue samples

Freshly dissected mouse liver tissue fragments were frozen in OCT (Thermo # 12678646) in liquid nitrogen cooled isopentane and stored at −80 °C. 10-μM-thick tissue sections were obtained after cryosection, mounted on Superfrost™ Plus Gold slides (Thermo Scientific, # K5800AMNZ72) and stored at −80 °C. Slides were thawed at RT, and rehydrated for 5 min in PBS then fixed in a 4% paraformaldehyde, PBS solution for 30 min at RT, washed 3 times in PBS, permeabilized for 30 min in a 1% a Triton X-100, PBS solution at RT, washed 3 times with PBS and saturated in a 0.5% BSA, PBS solution for 30 min at RT.

Incubation with primary antibodies was performed overnight in a 0.01% BSA, PBS solution at 4°C. Slices were washed 3 times with PBS and incubated with secondary antibodies for 2h in a 0.01% BSA, PBS solution at RT. Slices were incubated with 300nM DAPI (Invitrogen #D21490) in a PBS solution for 15 min at RT and in the dark and washed 3 times 5 min with PBS. Samples were mounted between the Superfrost™ Plus Gold slides and microscope cover glasses in ProLong Gold antifade reagent, left to polymerize overnight at RT and in the dark then sealed with nail polish. Antibody dilutions were the same as described for cells.

### Single molecule in situ hybridization immuno-fluorescence (smFISHIF) staining applied to tissue samples

Freshly dissected mouse liver tissue fragments were frozen in OCT in liquid nitrogen-cooled isopentane and stored at −80 °C. 10-μM-thick tissue sections were mounted on Superfrost™ Plus Gold slides (Thermo Scientific, # K5800AMNZ72) and stored at −80 °C. Slides were thawed at RT, and rehydrated for 5 min in PBS then fixed in a 4% paraformaldehyde, PBS solution for 30 min at RT, washed 3 times in PBS and permeabilized for 30 min in a 1% a Triton X-100, PBS solution at RT, washed 3 times in a 0.1% Tween 20 20 PBS solution, saturated in a 0.5% ultrapure BSA, 0.1% Tween 20, PBS solution for 30 min at RT and washed in a 10% formamide, 2X SSC solution for 20 min at RT. Overnight incubation and all following steps were performed as described for cells. Samples were mounted between the Superfrost™ Plus Gold slides and microscope cover glasses in ProLong Gold antifade reagent, left to polymerize overnight at RT and in the dark then sealed with nail polish. Antibody dilutions were the same as described for cells.

### Microscopy and Imaging

Immunofluorescence on mice liver slices was detected using an Axioscan 7 (Zeiss) equipped with a Set Orca hamamatsu Flash 4.0 V2 Axio Scan, using a dry 20x objective and controlled using Zen blue (version 3.7). IF and smFISHIF on cells and smFISHIF on liver slices were acquired using a Zeiss LSM980 confocal microscope (controlled with Zen blue 3.7) on an Airyscan 2 detector in Super Resolution mode with a 40X oil objective 1.3NA. GFP/Alexa-488 was excited using a 488 nm laser, Cy3/Alexa-555 were excited using a 561 nm laser, Alexa-670 was excited using a 633 nm laser. Figures were prepared with ImageJ (National Institutes of Health) and Illustrator (Adobe Systems).

### Fluorescence signal quantification

Fluorescence signal quantification in hollow hubs was done following these steps: i) confocal images were loaded in FIJI^84^ ii) a 3 mm line ROI was positioned to cross the hub in the larger region iii) Fluorescence intensity along the ROI was recovered at three Z planes (apical, mid and basal of the hubs) for the Ag and genome signals iv) intensity for the Ag and genome signals were plotted using Prism (GraphPad).

### RNA isolation and quantitative Reverse Transcription PCR (qRT-PCR)

Freshly dissected mouse liver tissue fragments were collected in 2mL tubes and flash frozen in liquid nitrogen. Total RNA were extracted using the QIAshredder (Qiagen, #79654) and Rneasy mini kit (Qiagen, # 74004) according to the manufacturer’s instructions. cDNA were synthetized from 1 mg of total RNA using the Maxima™ Minus cDNA Synthesis Master Mix kit (ThermoFisher, #M1662) according to the manufacturer’s instructions. qPCRs were performed using Power SYBR Green PCR Master Mix (Applied Biosystems, #4367659) on the CFX Opus 384 Real-Time PCR System (Bio-Rad, # 12011452). All of the primers used are provided in Supplementary Table 3. Data was analyzed using the CFX manager suite from Bio-Rad.

