## Supplementary Figures for "Unraveling human, rodent and snake *Kolmioviridae* replication to anticipate cross-species transmission"

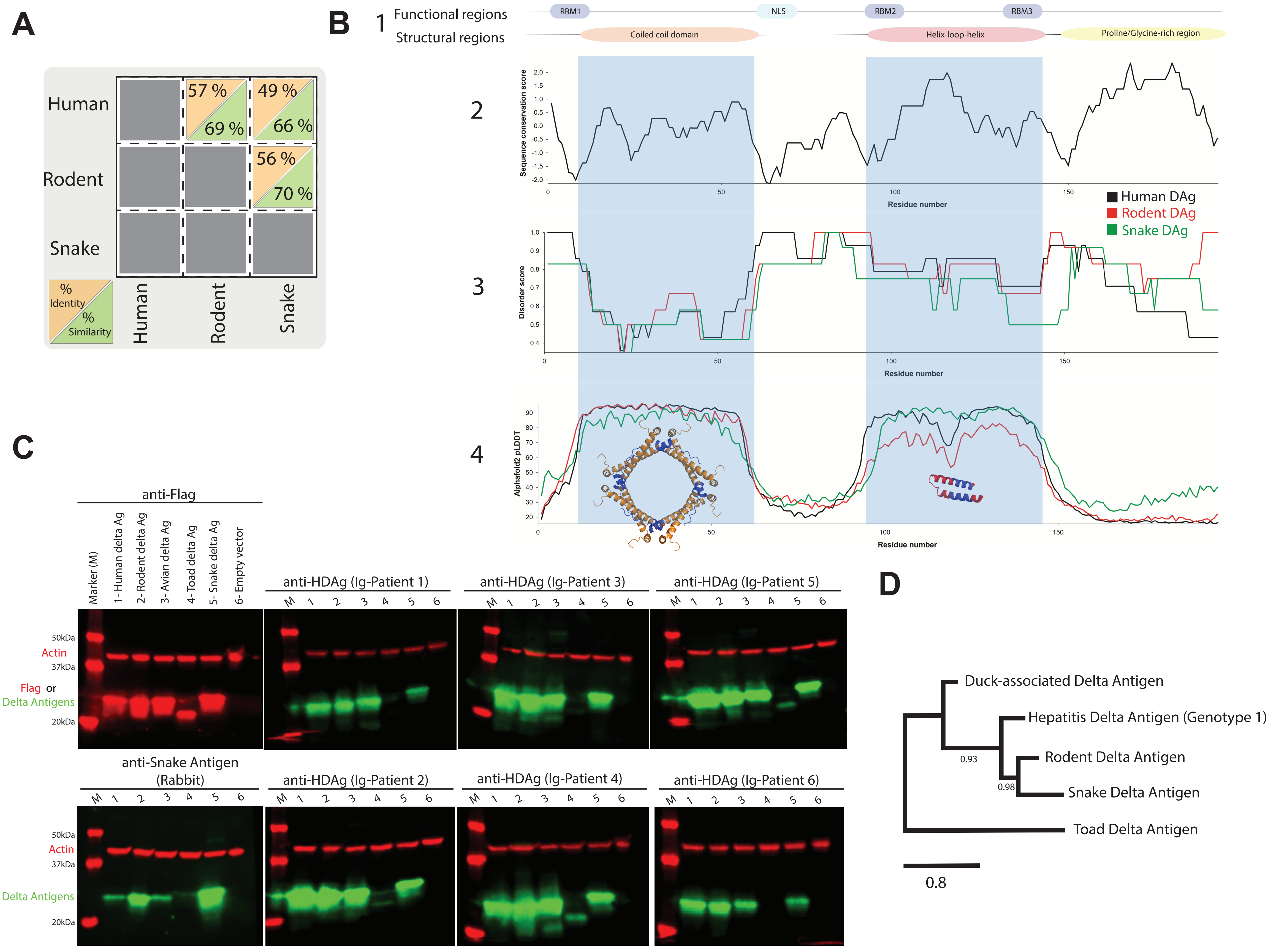


**Supplementary figure 1: DAgs homology and patient sera reactivity.** A-B) Tables recapitulating homologies between the full-length amino acid sequences of DAgs (A) and between regions corresponding to characterized HDV domains (B) 1. Structural and functional regions previously identified in HDAg. 2. Sequence conservation profile calculated from a sequence alignment of HDAg, RDAg and SDAg using AL2CO77 by applying a sliding average on a 10-residue window. 3. Predicted disorder propensity along the amino acid sequence for HDAg, RDAg and SDAg (shown as black, red and green lines, respectively). 4. Alphafold2 per-residue pLDDT (predicted local distance difference test) score versus residue number for the best scoring model of each DAg. The color code is the same as in 3. C) DAgs detection using purified patient sera. Huh7.5 cells were transfected with an empty pcDNA3.1 plasmid or with pcDNA3.1 plasmids encoding FLAG tagged DAgs from human (small), rodent, avian, toad, or snake deltaviruses. Cells were collected 3 days post-transfection and protein extracts were analyzed by western blot to detect DAgs expression and the presence of FLAG tags. Purified sera from 6 patients and from a rabbit immunized with SDAg were used. β-actin served as a loading control. D) Phylogenetic tree showing the relationships of HDAg (genotype 1), RDAg, SDAg, duck-associated DAg and toad DAg. Phylogenetic relationships were assessed by the maximum likelihood method available within PhyML (version 3.0)85.The significance of the branching order was estimated by the bootstrap method (1000 resampling). Only values >70% are shown.

Khalfi et al,_Supplementary Figure 2

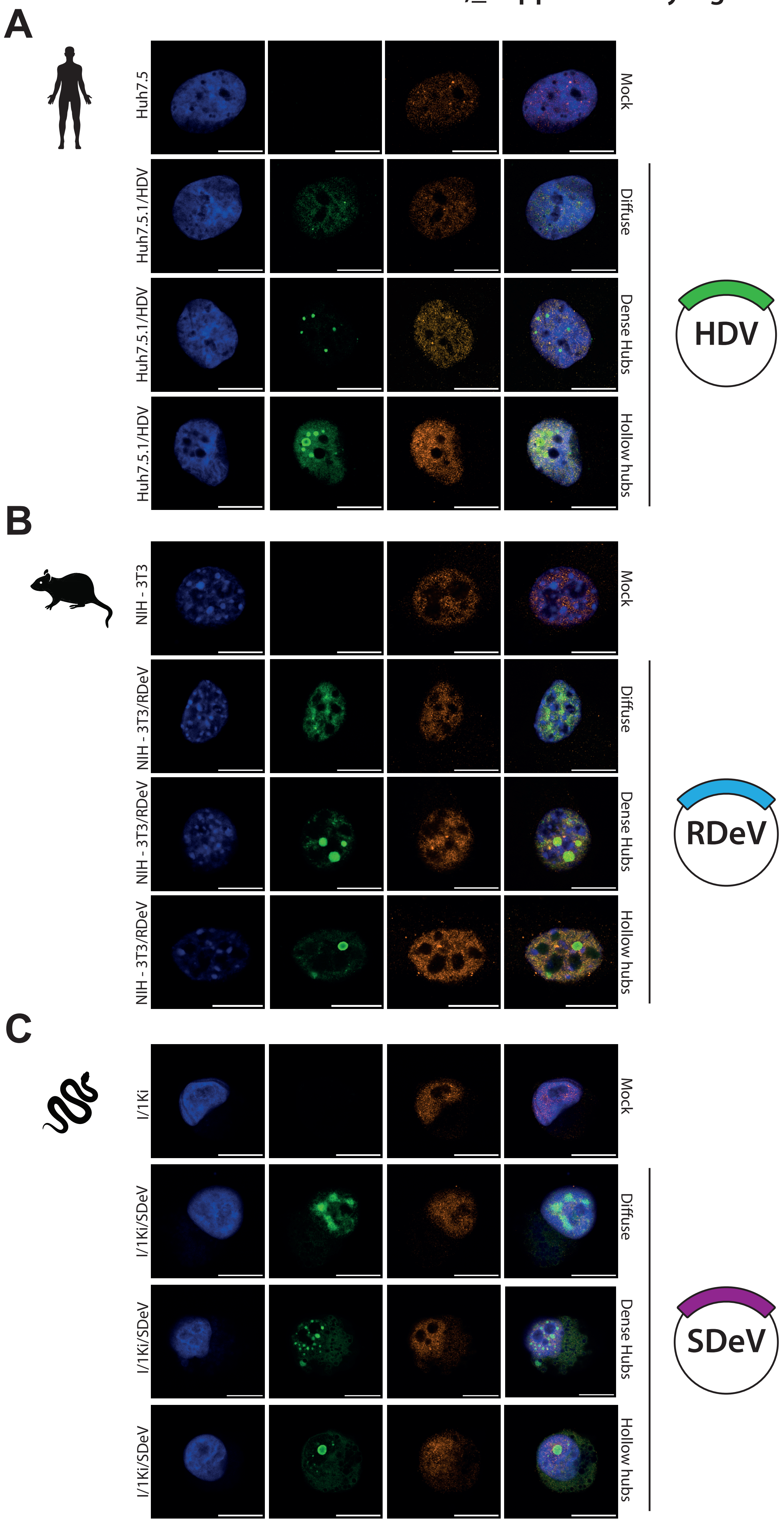


**Supplementary figure 2: RNA polymerase II localization in the nucleus of infected cells.** Huh7.5.1/HDV(A), NIH-3T3/RDeV (B) and I/1Ki/SDeV (C) cells were plated on microscopy slides and fixed to visualize DAgs and RNAPII localization in the nucleus. Corresponding uninfected cell lines served as negative controls. Representative confocal images are shown. Nuclei (in blue) were stained using DAPI and DAgs (in green) and RNAP II (in orange) were detected by IF, scale bars 10 µm. Cells were imaged on a LSM980 confocal microscope (Zeiss) and analyzed using ImageJ (version 2.9.0).

### Khalfi et al,_Supplementary Figure 2
